# Depside and depsidone synthesis in lichenized fungi comes into focus through a genome-wide comparison of the olivetoric and physodic acid chemotype of *Pseudevernia furfuracea*

**DOI:** 10.1101/2021.09.07.459332

**Authors:** Garima Singh, Daniele Armaleo, Francesco Dal Grande, Imke Schmitt

## Abstract

Primary biosynthetic enzymes involved in the synthesis of lichen polyphenolic compounds depsides and depsidones are Non-Reducing Polyketide Synthases (NR-PKSs), and cytochrome P450s (CytP450). However, for most depsides and depsidones the corresponding PKSs are unknown. Additionally, in non-lichenized fungi specific fatty acyl synthases (FASs) provide starters to the PKSs. Yet, the presence of such FASs in lichenized fungi remains to be investigated. Here we implement comparative genomics and metatranscriptomics to identify the most likely PKS and FASs for the synthesis of olivetoric and physodic acid, the primary depside and depsidone defining the two chemotypes of the lichen *Pseudevernia furfuracea*. We propose that the gene cluster PF33-1_006185, found in both chemotypes, is the most likely candidate for olivetoric and physodic acid biosynthesis. This is the first study to identify the gene cluster and the FAS likely responsible for physodic and olivetoric acid biosynthesis in a lichenized fungus. Our findings suggest that gene regulation and other epigenetic factors determine whether the mycobiont produces the depside or the depsidone, providing the first direct indication that chemotype diversity in lichens can arise through regulatory and not only through genetic diversity. Combining these results and existing literature, we propose a detailed scheme for depside/depsidone synthesis.

## Introduction

Depside and depsidones, the polyphenolic polyketides mostly synthesized by lichenized fungi, are of significant pharmaceutical interest (Shukla *et al*., 2010; Shrestha and St. Clair, 2013; Ingelfinger *et al*., 2020). Depsides consist of two or sometimes three orcinol or ß-orcinol-derived aromatic rings joined by ester linkages; depsidones have an additional ether linkage between the rings (Fig. 1). Additionally, depending on the starters used by the polyketide synthases (PKSs) assembling their backbones, 3-7 carbon side chains may be linked to the 6 and 6’ carbons of the orcinol-derived rings. Together with other ring modifications, side chains constitute the distinguishing features of different depsides and depsidones. Although chemical proposals for depside and depsidone biosynthesis go back many decades (Seshadri, 1944; Elix *et al*., 1987), the precise enzymatic steps of depside and depsidone synthesis still need to be elucidated. Furthermore, for most of the depside and depsidone metabolites of lichens the corresponding genes remain uncharacterized. This is because fungi contain far more biosynthetic genes than known compounds (Meiser *et al*., 2017; Calchera *et al*., 2019). One way to connect metabolites to the associated genes is to identify all the genomic regions with putative biosynthetic genes, and narrow down this selection to the most likely gene cluster based on phylogenetic evidence, and other cluster information, such as presence of particular genes. Long reads sequencing technologies providing high quality contiguous genome assemblies have greatly facilitated this process.

**Figure 1.**
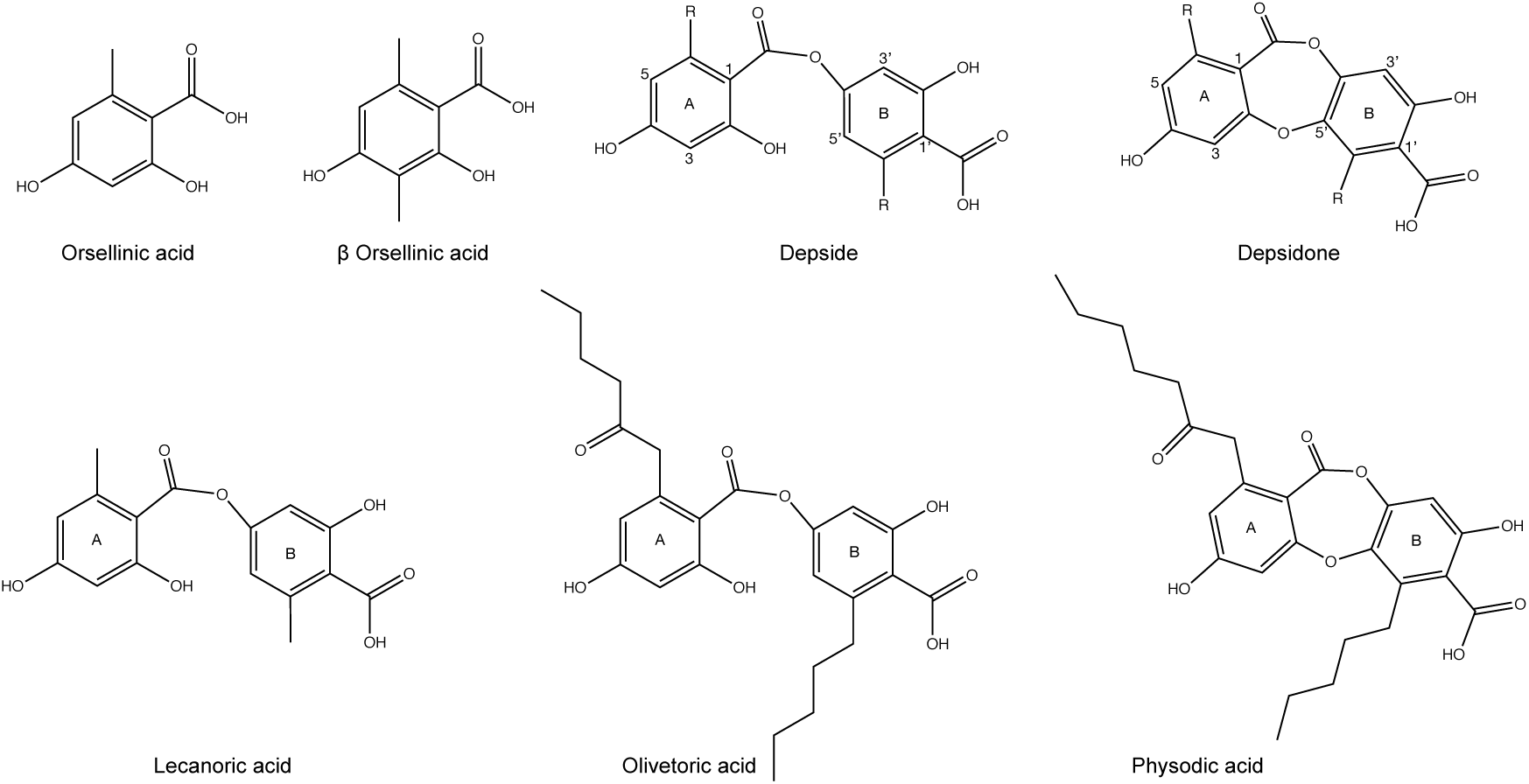
Chemical structure of orsellinic acid and methyl-3-orsellinate (the monocyclic precursors of depsides and depsidones), lecanoric acid, a depside, a depsidone, olivetoric acid, and physodic acid. The letters and ring numberings used for the generic depside are the same for all depsides and depsidones.

Fungal type I PKSs are iterative, and consist of several domains with defined functions. Non-reducing, type I PKSs (NR-PKSs) contain KS (keto-synthase), AT (acyltransferase), PT (product template), ACP (acyl carrier protein), and TE (thioesterase) domains (Kroken *et al*., 2003; Cox and Simpson, 2009). While NR-PKSs have long been known to assemble and fold the carbon backbones of depsides and depsidones, their specific roles in linking the rings have come to light only recently. A single PKS catalyzes the formation and dimerization of phenolic rings to produce a depside (Armaleo et al, 2011; Kealey et al 2021) while a cytochrome P450 is needed to catalyze the formation of an ether bond between the depside rings to produce a depsidone (Armaleo *et al*., 2011). The PKS constructs and esterifies the two different rings using two ACP domains (Feng *et al*., 2019). The genes for PKS and cytochrome P450 (CytP450) are closely linked within the same biosynthetic gene cluster (BGC). BCGs contain several genes involved in the synthesis of a compound, e.g., the core biosynthetic PKS, redox enzymes, transporters, etc. A BGC is named based on the type of backbone enzyme encoded by the core gene, e.g., a PKS cluster, a terpene cluster, etc.

Apart from the above-stated genes, studies with non-lichen-forming fungi indicate that specific fatty acyl synthases (metabolite FASs) play significant roles in metabolite synthesis by providing the appropriate acyl chain starters to some PKSs (Hitchman *et al*., 2001; Watanabe and Townsend, 2002; Smith and Tsai, 2007). Inactivation of metabolite FASs may inhibit secondary metabolite synthesis even when the corresponding PKS and BGC remain functional (Brown *et al*., 1996). However, the role of metabolite FAS in providing the starters for depsides and depsidones in lichen-forming fungi has not been investigated, despite the fact that the primary starters of orcinol depsides and depsidones are, besides the C2 doublet from AcetylCoA, C4, C6, and C8 acyl chains (Culberson and Culberson, 1976). After polyketide assembly and cyclization, the resulting ring sidechains will respectively be 1, 3, 5, 7 carbons long. Recently and coincidentally, the NR-PKS from the lichen *Pseudevernia furfuracea* that we identify in this work as the likely producer of the depside olivetoric acid was reported to produce the depside lecanoric acid when heterologously expressed in yeast (Kealey *et al*., 2021). Yet in nature lecanoric acid has never been reported from *P. furfuracea*. The difference between lecanoric and olivetoric acid is that the former has a methyl group on each ring whereas the latter has as a C5 side chain on one ring and a C7 side chain on the other (Fig. 1). We integrate this apparent discrepancy with other data to highlight the central role that lichen short chain FASs are likely to play in providing the side chains common in orcinol depsides and depsidones.

In this study we implemented a long-read-based genomic approach to better understand the mechanism of depside/depsidone synthesis in lichen-forming fungi. We chose *Pseudevernia furfuracea* as our study system, because it is a textbook example of chemosyndrome variation in a lichen-forming fungus (Culberson *et al*., 1977; Halvorsen and Bendiksen, 1982; Kosanić *et al*., 2013). This lichen consists of two naturally occurring chemotypes, one synthesizing an orcinol depside (olivetoric acid), and the other the corresponding depsidone (physodic acid), and thus constitutes an ideal model to study depside/depsidone synthesis, and the causes of chemotype diversity. Both chemotypes also synthesize the ß-orcinol depside atranorin, common in many lichens. Specifically, we aim to answer the following questions: 1) Do the depside and the depsidone producer contain the same number of BGCs? 2) Which BGC/s are likely responsible for the production of depside olivetoric acid and the depsidone physodic acid in *P. furfuracea* chemotypes? 3) Are there homologs of metabolite FASs in the lichen-forming fungal genome? 4) Can we integrate the available data to provide a detailed scheme of orcinol depside/depsidone biosynthesis in lichens?

## Materials and methods

### Identification of chemotypes

We used high performance liquid chromatography (HPLC) to investigate the chemotype of *P. furfuracea*. For this, we collected several samples of *P. furfuracea* and performed HPLC analysis using the protocol from Feige et al. and Benatti et al. (2013). Firstly, small thallus pieces were extracted for 1 hour at room temperature in 200 µl of methanol. From this, 150 µl of the extract of each sample was centrifuged 1 min at 800 rpm through a Pall Acroprep Advance 0.2 µm polytetrafluoroethylene filter plate and then diluted 10-fold with methanol. The samples were analyzed on an Agilent 1260 quaternary system with a quaternary pump, an incorporated degasser and using an Agilent Poroshell 120 EC-C18 column (2.7 µm, 3.0 × 50 mm). Substances were separated at 30°C using two solvent systems and a flow rate of 1.4 ml/min. Solvent A is Aqua Bidest, 30% methanol and 0.0658% trifluoroacetic acid, and solvent B is 100% methanol. The HPLC system was equilibrated to solvent A for 2 min and 2µl of extract was injected automatically after a needle wash. The runs continued isocratically for 0.18 min, solvent B was increased to 58% within 5 min, then increased to 100% within the next 5 min and isocratically maintained for 0.82 min. The runs ended with solvent A being increased back to 100% within 0.5 min. After the run the column was flushed for two minutes before the next run. Compounds were detected with a diode array detector (DAD) at 210, 254, 280 and 310 nm. The retention times and spectra (λ = 190-650 nm with 2 nm steps) were compared against a library of authentic products derived under identical conditions using the Agilent OpenLAB CDS ChemStation software. We then selected one sample of each chemotype for genome sequencing (Supplemenraty Table S1).

### DNA extraction and genome sequencing

Lichen thalli were thoroughly washed with sterile water, and checked under the stereomicroscope for the presence of possible contamination. DNA was extracted from both samples using a CTAB-based method (Cubero and Crespo, 2002). DNA concentration was measured with a Qubit fluorometer (dsDNA BR, Invitrogen). 4.1 µg and 7.4 µg DNA for the physodic- and olivetoric acid chemotype, respectively, were sent to Novogene Hong Kong for PacBio library preparation and sequencing on two separate SMRT cells, one for each chemotype.

### Genome assembly and annotation

PacBio metagenomes were assembled using the long-read based assembler metaFlye v2.3.1 (Kolmogorov *et al*., 2019). Reads were filtered for length (>2000 kb fragments only) and assembly was optimized for minimal read overlap of 3 kb, and an estimated combined metagenome size of 120 Mb. The assembled genome was polished twice using the software Arrow from the SMRTlink suite v. 5.0.1.9585 (Walker *et al*., 2014). The resulting contigs were then scaffolded with SSPACE-LongRead v1.1 (Boetzer and Pirovano, 2014). Ascomycota contigs were then identified in the metagenomic assembly using Diamond v0.8.34.96 BLASTx using the more-sensitive mode for longer sequences and a default e-value cut-off of 0.001 against the custom database. The Diamond results were then parsed in MEGAN68 v.6.7.7 using max expected set to 1E-10 and the weighted lowest common ancestor (LCA) algorithm. All contigs assigned to Ascomycota were exported to represent the *P. furfuracea* mycobiont. Assembly indicators such as number of contigs, total length and N50 were accessed with Assemblathon v2 (Table 1). Genome completeness was estimated based on evolutionarily-informed expectations of gene content with BUSCO v.4.0 (Benchmarking Universal Single-Copy Orthologs) (Simão *et al*., 2015). The genomes are deposited in GenBank under accessions xx and xx.

**Table 1.**
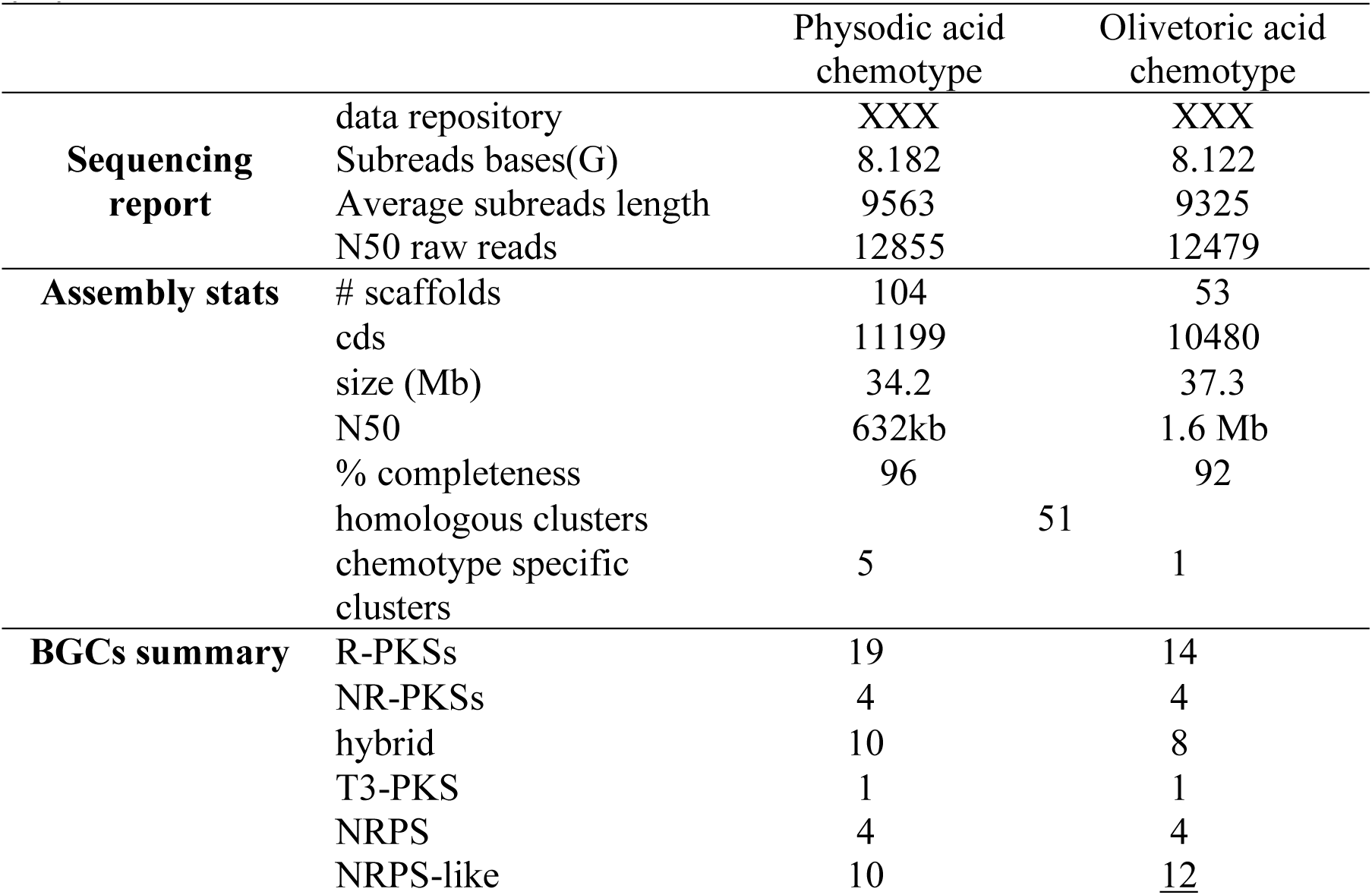
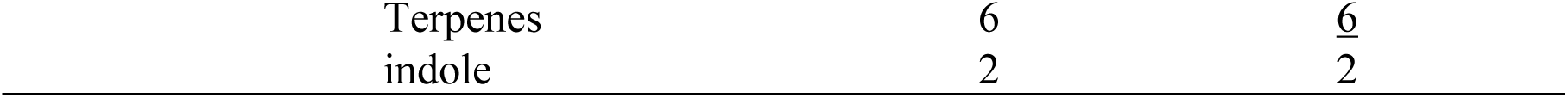
Genome statistics and GB accession numbers of the two chemotypes of *P. furfuracea*. Biosynthetic gene clusters (BGCs) predicted by antiSMASH are also given. PKS=polyketide synthase, R-PKS=reducing PKS, NR-PKS=non-reducing PKS T3 PKS=xx, hybrid=xx, NRPS=non-ribosomal peptide synthetase.

### Identification and Annotations of Biosynthetic Gene Clusters

Gene prediction, functional annotation and prediction of BGCs in both *P. furfuracea* chemotype assemblies were performed with scripts based on the funnannotate pipeline (Palmer and Stajich, 2019) and antiSMASH (antibiotics & SM Analysis Shell, v5.0) (Medema *et al*., 2011; Blin *et al*., 2019). First, the repetitive elements were masked in the assembled genomes (using funannotate), followed by gene prediction using BUSCO2 to train Augustus and self-training GeneMark-ES. Functional annotation was then automatically carried out with InterProScan, Eggnog-mapper and BUSCO ascomycota_odb10 models. Secreted proteins were predicted using SignalP as implemented in funannotate ‘annotate’ command. The interproscan, antismash and phobius results were automatically generated.

### Identification of homologous BGCs

Homologous clusters between the two *P. furfuracea* chemotypes were identified by performing reciprocal blast between the core genes of the BGCs of both genomes. For this, first the core genes from the predicted BGCs of one chemotype were used as database and the core genes of the BGCs from the other chemotype as query. The process was then repeated using the other chemotype as database. The homology between the clusters was then confirmed based on sequence similarity and the most similar hit of the core gene in the MIBiG v2 (Minimum Information about a Biosynthetic Gene cluster; (Kautsar *et al*., 2020)) database (Supplementary table S2).

Homologous clusters were visualized using synteny plots as implemented in Easyfig v2.2.3 (Sullivan *et al*., 2011). The GBK input files for Easyfig were generated with seqkit v0.10.1(Shen *et al*., 2016) and the seqret tool from EMBOSS v6.6.0.0 (Rice *et al*., 2000). Easyfig was run with tblastx v2.6.0+, a minimum identity value of 90 and a minimum length of 50 to draw the blast hits (Kjærbølling *et al*., 2018). Clusters were manually matched for orientation so that the core gene were oriented in the same direction. For six BGCs, no corresponding cluster was detected in the other chemotype (Table 2, Supplementary Table S2).

**Table 2.**
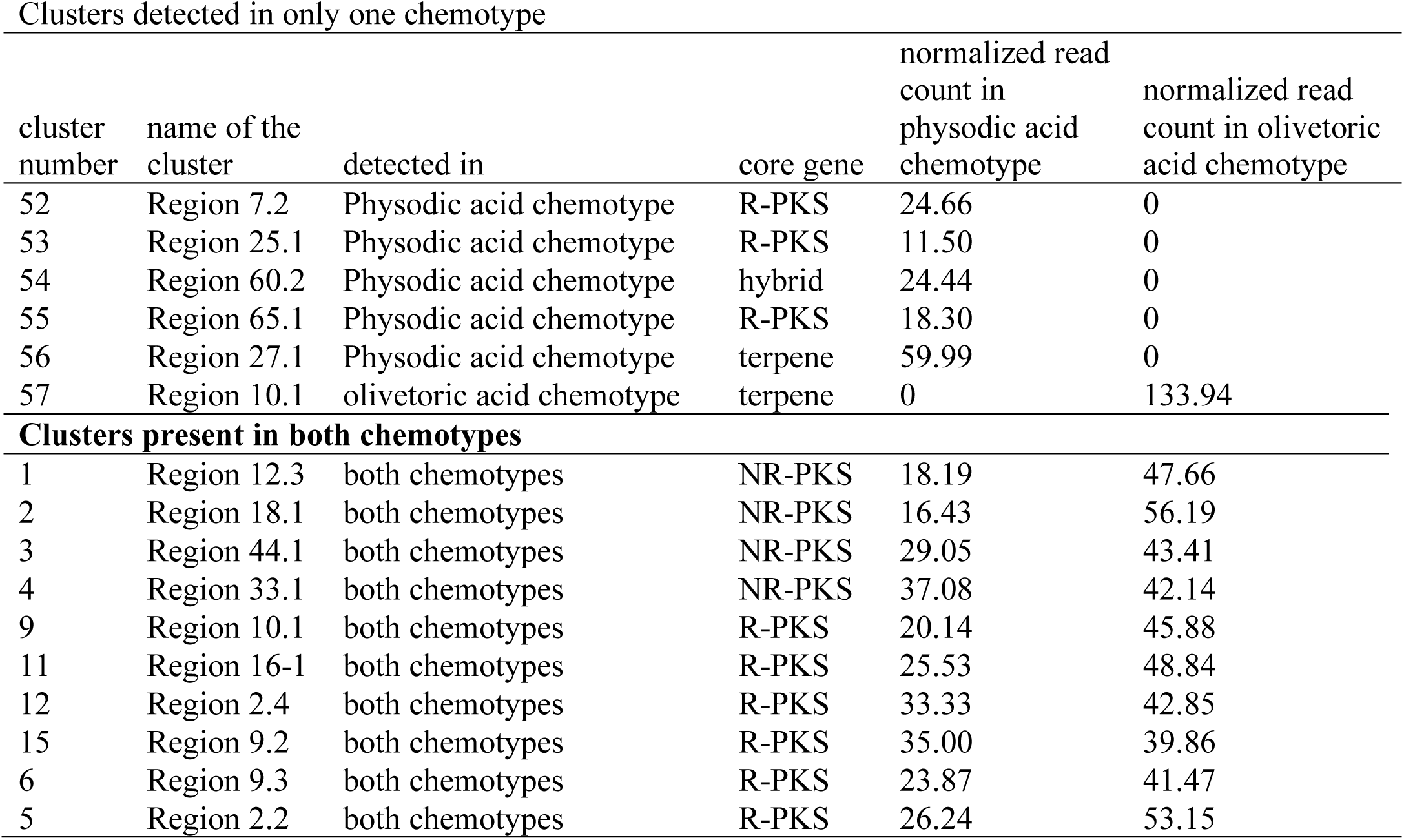
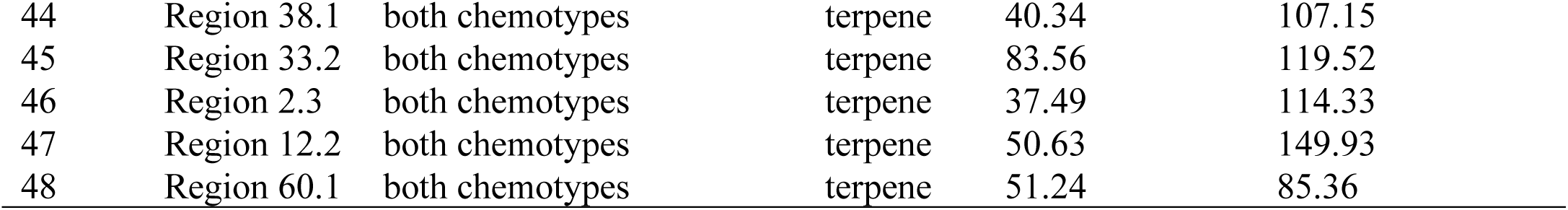
Number of raw DNA reads (nucleotides) normalized by number of reads and gene length aligned to the core genes of the clusters detected in only one chemotype Properties of the clusters detected in only one chemotype including the core gene and its length, number of raw reads (nucleotide) normalized read counts (by number of reads and gene length (RPKM approach)) aligned to the core genes of the clusters detected in only one chemotype.

| Clusters detected in only one chemotype |  |  |  | normalized read count in physodic acid chemotype | normalized read count in olivetoric acid chemotype |
| --- | --- | --- | --- | --- | --- |
| cluster number | name of the cluster | detected in | core gene |  |  |
| 52 | Region 7.2 | Physodic acid chemotype | R-PKS | 24.66 | 0 |
| 53 | Region 25.1 | Physodic acid chemotype | R-PKS | 11.50 | 0 |
| 54 | Region 60.2 | Physodic acid chemotype | hybrid | 24.44 | 0 |
| 55 | Region 65.1 | Physodic acid chemotype | R-PKS | 18.30 | 0 |
| 56 | Region 27.1 | Physodic acid chemotype | terpene | 59.99 | 0 |
| 57 | Region 10.1 | olivetoric acid chemotype | terpene | 0 | 133.94 |
| Clusters present in both chemotypes |  |  |  |  |  |
| 1 | Region 12.3 | both chemotypes | NR-PKS | 18.19 | 47.66 |
| 2 | Region 18.1 | both chemotypes | NR-PKS | 16.43 | 56.19 |
| 3 | Region 44.1 | both chemotypes | NR-PKS | 29.05 | 43.41 |
| 4 | Region 33.1 | both chemotypes | NR-PKS | 37.08 | 42.14 |
| 9 | Region 10.1 | both chemotypes | R-PKS | 20.14 | 45.88 |
| 11 | Region 16-1 | both chemotypes | R-PKS | 25.53 | 48.84 |
| 12 | Region 2.4 | both chemotypes | R-PKS | 33.33 | 42.85 |
| 15 | Region 9.2 | both chemotypes | R-PKS | 35.00 | 39.86 |
| 6 | Region 9.3 | both chemotypes | R-PKS | 23.87 | 41.47 |
| 5 | Region 2.2 | both chemotypes | R-PKS | 26.24 | 53.15 |
| 44 | Region 38.1 | both chemotypes | terpene | 40.34 | 107.15 |
| 45 | Region 33.2 | both chemotypes | terpene | 83.56 | 119.52 |
| 46 | Region 2.3 | both chemotypes | terpene | 37.49 | 114.33 |
| 47 | Region 12.2 | both chemotypes | terpene | 50.63 | 149.93 |
| 48 | Region 60.1 | both chemotypes | terpene | 51.24 | 85.36 |

### Phylogenetic analyses

NR-PKSs have been divided into nine groups based on protein sequence similarity and PKS domain architecture (Ahuja *et al*., 2012; Liu *et al*., 2015; Kim *et al*., 2021). We took representative PKSs from each group (amino acid sequences) and added the amino acid sequences of the eight *NR-PKSs* from *P. furfuracea*. The dataset includes 107 PKS sequences from *Cladonia borealis, C. grayi, C. macilenta, C. metacorallifera, C. rangiferina, C. uncialis, Pseudevernia furfuracea* and *Stereocaulon alpinum*. Sequences were aligned using MAFFT as implemented in Geneious v5.4. Gaps were treated as missing data. The maximum likelihood search was performed on the aligned amino acid sequences with RAxML-HPC BlackBox v8.1.11 (Stamatakis, 2006, 2014) on the Cipres Scientific gateway (Miller *et al*., 2010).

### Candidate cluster for physodic- and olivetoric acid synthesis

In addition to the phylogenetic evidence, we implemented several criteria to select the candidate cluster for depside/depsidone synthesis in *P. furfuracea*: 1) it must be present in both chemotypes, (presence in both chemotypes is expected as the basic structure of physodic and olivetoric acid is same except that physodic acid contains an additional ether bond (Fig. 1)) 2|) it must contain a *NR-PKS* (the non-reduced backbone of lichen depsides/depsidones suggests that the *PKSs* involved in their synthesis are *NR-PKSs*), and 3) the *NR-PKS* must contain two ACPs (the presence of two ACPs has been associated with depside production in fungi (Feng *et al*., 2019; Lünne *et al*., 2020), and is a typical feature of lichen-forming fungal *NR-PKSs* involved in depside/depsidone synthesis (Armaleo *et al*., 2011; Pizarro *et al*., 2020)). Additionally, in the physodic acid producer the candidate BGC must contain a *CytP450* which produces depsidones by forming the ether bond between the two orsellinic rings of the depside (Armaleo *et al*., 2011).

Summarizing, the following criteria were used for the identification of olivetoric-/physodic acid BGC: 1) the candidate BGC should be homologous and present in both chemotypes, 2) presence of a *CytP450*, and 3) presence of two ACP domains in the *PKS* as the ring dimerization of orsellinic acid precursors into a depside involves two ACP domains (Feng *et al*., 2019; Lünne *et al*., 2020).

### Identification of HexA and HexB

Metabolite FASs consist of a HexA/HexB multienzyme complex (Brown *et al*., 1996; Hitchman *et al*., 2001). Homologous of *HexA* and *HexB* were identified by blasting (blastN) the H*exA* and *HexB* homologs of *Cladonia grayi* (CLAGR_008938-RA and CLAGR_008939-RA, available at https://mycocosm.jgi.doe.gov/cgi-bin/browserLoad/?db=Clagr3&position=scaffold_00085:34887-93856) against the genomes of both chemotypes.

### Metatranscriptome analyses and quantification of *PKS, CytP450* and *HexA* and *HexB* transcripts

The details of RNA isolation and transcriptome extraction are given in Meiser et al. (2017). Briefly, for RNA isolation, whole lichen thalli were collected and stored directly in RNAlater (Sigma-Aldrich Chemie GmbH, Munich, Germany). RNA was isolated from both chemotypes of *P. furfuracea* by using the method described by Rubio-Piña & Zapata-Pérez (2011) after blotting the thalli dry and grinding them in liquid nitrogen with a mortar and pestle. The isolated poly-A+ RNA was further purified with the RNeasy MinElute Clean-up Kit (Qiagen, Hilden, Germany), and sequenced (250 bp paired-end reads) on Illumina MiSeq at StarSeq (Mainz, Germany).

The BGC for depside/depsidone biosynthesis in each chemotype contains 10 genes including one *Pfur33-1_006185*, one *CytP450*, and a monooxygenase (see below). The other seven code for unidentified proteins. We used transcriptome data to check which genes in this cluster are transcriptionally active. For this, we first indexed the sequence of interest using bowtie and then aligned it to the transcripts (both paired and unpaired reads) using tophat v2 (Kim *et al*., 2013). To make the counts comparable between chemotypes we used RPKM normalization of the read counts(Mortazavi *et al*., 2008), accounting for sequencing depth and gene length (Oshlack and Wakefield, 2009; Robinson and Oshlack, 2010; Dillies *et al*., 2013). The normalization for sequencing depth was performed by dividing the counts by the raw read count of the given gene with the total number of reads in each sample. The resulting number was then divided by gene length in kilobases to obtain RPKM normalized counts.

## Results

### Genomes of the *P. furfuracea* chemotypes

The number of reads for each sample retained after quality and length filtering is given in Table 1. The reference genome of the PacBio-based *P. furfuracea* physodic acid chemotype (NCBI acc. no. XXX) is ∼34 Mb in length and has a completeness of 96% according to BUSCO (details in Table 1). The genome of the olivetoric acid chemotype (NCBI acc. no. XXX) is ∼37 Mb in length and has a completeness of 92% according to BUSCO (details in Table 1).

### Predicted BGCs

A total of 51 homologous BGCs were present in both chemotypes: 14 clusters with reducing *PKSs* (*R-PKS*), eight clusters with *NR-PKSs*, one cluster with type III *PKS*, seven hybrid clusters, 14 clusters with *NRPS* or *NRPS*-like genes, five clusters with terpene synthase, and two clusters with indole synthase as a core gene (Supplementary Table S2). Six BGCs, were found only in one of the two chemotypes (Table 2). Five BGCs were only present in the physodic acid chemotype (four BGCs with a *R-PKS*, a hybrid cluster with a *R-PKS* and a *NRPS*, and a cluster with terpene synthase as the core gene), and one BGC with terpene synthase as the core gene was present only in the olivetoric acid chemotype (Table 2).

### Phylogenetic analyses

Out of eight *NR-PKSs*, only two, *Pfur33-1_006185* and *Pfur2-2_003072*, grouped into phylogenetic group I, whose *PKSs* are involved in the synthesis of orcinol derivatives (Fig. 2), like grayanic, olivetoric and physodic acid. Of these *PKSs, Pfur33-1_006185* was closely related to *PKS16*, the *PKS* associated with grayanic acid biosynthesis from *Cladonia grayi*, whereas *Pfur2-2_003072* was closely related to *PKS27*. The Pfur33-1_006185 cluster also contains a *CytP450* next to the *NR-PKS* in an arrangement analogous to that in the PKS16 cluster in *C. grayi*.

**Figure 2.**
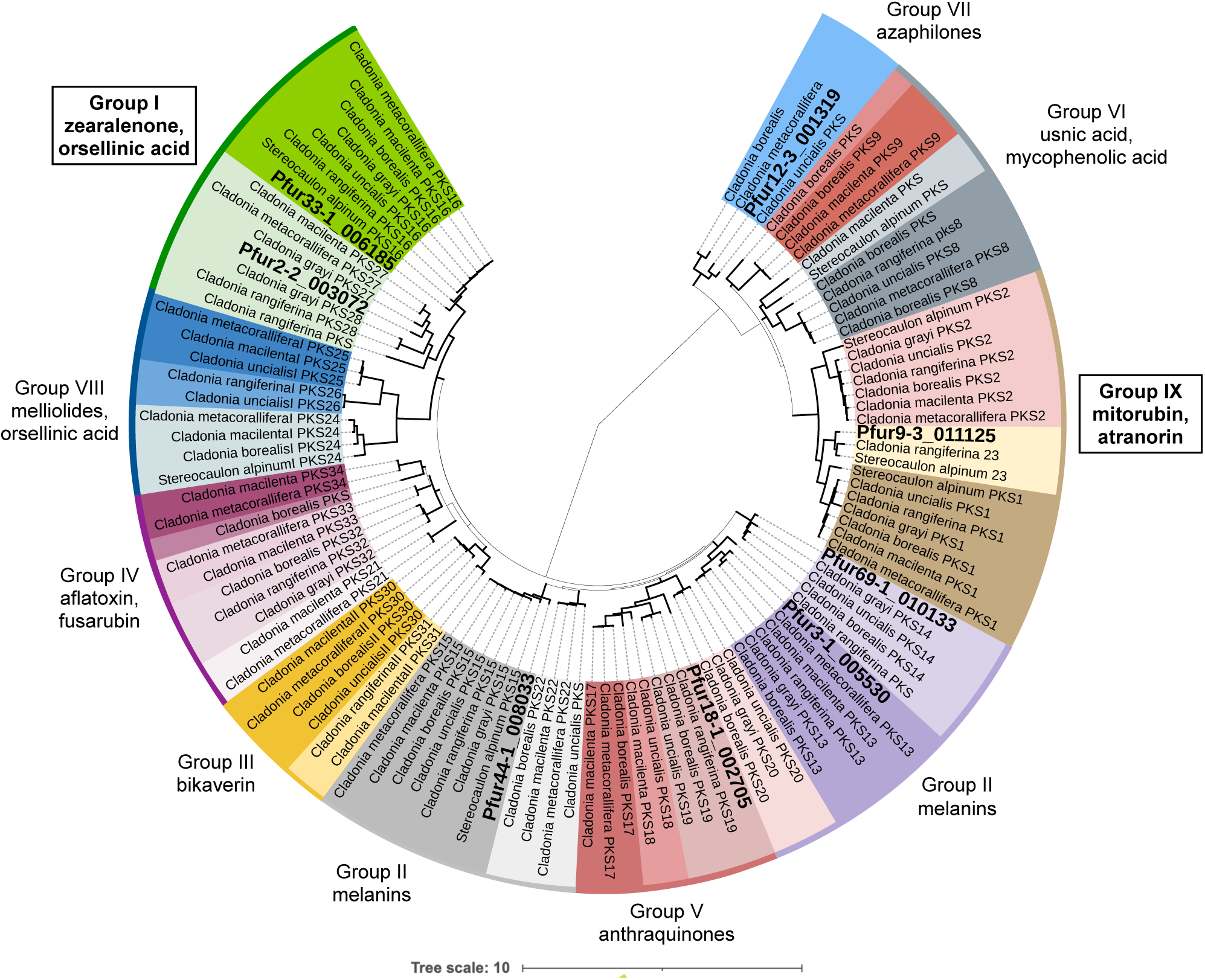
A maximum likelihood tree of 107 NR-PKSs amino acid sequences from six *Cladonia* spp., *Stereocaulon alpinum* and eight *P. furfuracea*. Branches is bold indicate the bootstrap support of >70%. Different colors indicate different PKSs. PKS families are based on Kim et al. (2021). PKSs of *P. furfuracea* are indicated in bold and with larger fonts.

### Selection of the candidate cluster for physodic acid and olivetoric acid

We complemented the phylogenetic evidence with the other characteristics of the cluster to select the possible olivetoric/physodic acid cluster. We found eight clusters containing a *NR-PKS* that were present in both chemotypes (Supplementary Table 2, Table 3). Of these, only one cluster, cluster 4 (Table 3), contained a *NR-PKS* with two ACP domains and a *CytP450* in the cluster. The domains of this PKS are – SAT-KS-AT-ACP-ACP-TE (Fig. 3). The most similar *PKS* to this is the *PKS* linked to grayanic acid biosynthesis, PKS16. We therefore propose cluster 4 to be the most likely candidate for olivetoric-/physodic acid biosynthesis in *P. furfuracea*. The cluster has an almost identical structure in both chemotypes, with 10 genes including a *NR-PKS*, a *CytP450* and a monooxygenase (Fig. 3). The protein products of the remaining seven genes are unidentified.

**Table 3.** NR-PKS clusters detected in both chemotypes of *P. furfuracea*, the PKS domains and other genes present in the cluster Properties of the *NR-PKS* clusters detected in both the chemotypes of *P. furfuracea*. NR-PKS= non-reducing PKS. The cluster in bold, in the box (cluster 4 containing *Pfur33-1_006185*), is the likely cluster for depside/depsidone synthesis. The domain acronyms stand for: KS = keto-synthase, AT=acyltransferase, ACP=acyl carrier protein and KR= ketoreductase. The PKS category is based on the phylogenetic placements of the *P. furfuracea* NR-PKSs in the PKS groups from (Kim *et al*., 2021). Cluster number refers to Supplementary table S2 and the PKS number to the antiSMASH cluster, followed by the gene number.

| cluster number | PKS | compound defining the group (Kim et al 2021) | PKS category (Kim et al 2021) | PKS domains | total genes in the cluster | identified genes in cluster | Most similar known PKS (MIBiG) | coverage & similarity to MIBiG cluster |
| --- | --- | --- | --- | --- | --- | --- | --- | --- |
| 1 | Pfur12-3_001319 | VII (azaphilones, monascorubrin) | not known | SAT-KS-AT-ACP cMT-TD | 12 | Regulatory gene, NRPKS | monascorubrin | 95%, 58% |
| 2 | Pfur18-1_002705 | V (anthraquinones) | PKS20 | SAT-KS-AT-PT-ACP | 16 | metallo-beta-lactamase family protein, NRPKS, halogenase | RES-1214-2 | 61%, 100% |
| 3 | Pfur44-1_008033 | II (melanins) | PKS15 | SAT-KS-AT-PT-ACP ACP-TE | 11 | PKS, dehydrogenase/reductase (KR) | naphthalene | 99.5%, 53% |
| **4** | **Pfur33-1_006185** | **I (zearalenone, orsellinic acid)** | **PKS16** | **SAT-KS-AT-ACP ACP-TE** | **10** | **cyt P450, PKS, monooxygenase** | **grayanic acid** | **99%, 73%** |
| 5 | Pfur2-2_003072 | I (zearalenone, orsellinic acid) | PKS27 | SAT-KS-AT-PT-ACP-TE | 13 | Omethyltransferase, cyt P450, crotonyl-CoA reductase / alcohol dehydrogenase, red-PKS, NRPKS, GATase 7 | grayanic acid | 100%, 38% |
| 6 | Pfur9-3_011125 | IX (mitorubin, atranorin) | PKS23 | SAT-KS-AT-PT-ACP cMT | 11 | alkyl hydroperoxide reductase/ Thiol specific, PKS, cyt P450, drug resistance transporter | ascochlorin cluster (*Acremonium egyptiacum*) | 100%, 38% |
| 7 | Pfur3-1_005530 | II (melanins) | PKS13 | SAT-KS-AT-PT-ACP-ACP-TE | 13 | serine/threonine protein kinase, Drug resistance transporter, monooxygenase FAD-binding, NRPKS, short-chain dehydrogenase/reductase SDR, O-methyltransferase, transcription regulator |  | 100%, 48% |
| 8 | Pfur69-1_010133 | II (melanins) | PKS14 | SAT-KS-AT-PT-ACP-ACP-TE | 9 | Drug resistance transporter, O-methyltransferase, halogenase, monooxygenase FAD-binding, adh_short (DH-KR), NRPKS, monooxygenase FAD-binding | 6-hydroxymellein (*Cladonia uncialis*) | 99%, 78% |

**Figure 3.**
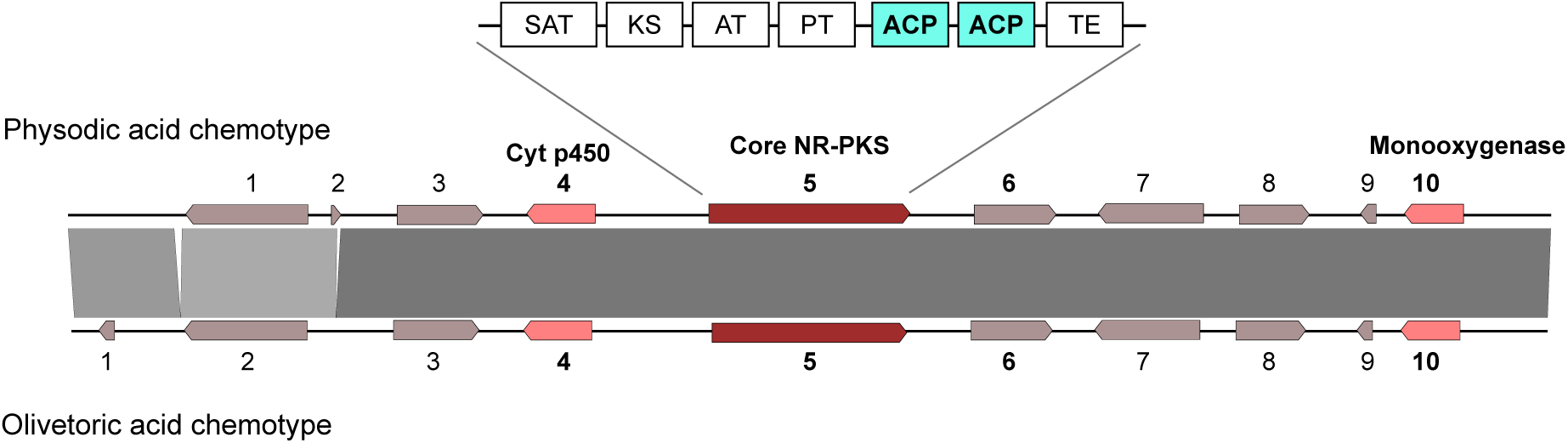
Synteny plot based on tBLASTn depicting conservation and synteny between the homologous putative cluster for the depsidone, physodic acid and the depside, olivetoric acid synthesis in *P. furfuracea*. Bolded numbers represent the *P. furfuracea* PKSs.

We excluded the other seven NR-PKS clusters as possible candidates for olivetoric/physodic acid synthesis based on the following reasons: Cluster 1 has a cMT domain in the PKS, lacks *CytP450* and it *PKS* groups with group VII PKSs associated with the synthesis of azaphilones, monascorubrin related compounds. The *PKS* of cluster 2 lacks a TE domain, and it groups with *PKSs* linked to anthraquinone biosynthesis (group VII). *Cytochrome P450* is not present in this cluster. In cluster 3 *CytP450* is absent and the *PKS* groups with PKSs associated with melanin synthesis (group V). Cluster 5 contains a *R-PKS* and a *NR-PKS* next to each other divergently transcribed from the same region, suggesting common regulation and no connection to olivetoric acid synthesis. This cluster does contain a *CytP450*, but also an *O*-methyltransferase (*OMT*) which is not required for olivetoric/physodic synthesis. Cluster 6 contains a *CytP450*, but the *PKS* has a cMT domain and groups phylogenetically with atranorin-synthesizing PKSs (group IX). This is the most likely cluster involved in the synthesis of atranorin (see next paragraph). Clusters 7 and 8 lack *CytP450*, contain an OMT, and the PKSs group with melanin synthesizing PKSs (group II).

### A putative atranorin cluster is present in *P. furfuracea*

Apart from the orcinol-derived olivetoric- and physodic acid, atranorin (a ß-orcinol depside) is a common secondary metabolite produced by both chemotypes of *P. furfuracea*. Recently, the atranorin cluster from *Cladonia rangiferina* was characterized and heterologously expressed (Kim *et al*., 2021). The atranorin PKS, PKS23, belongs to group IX (Fig. 2). The putative atranorin PKS is expected to have the following domains: SAT-KS-AT-PT-ACP-cMT-TE. In our study, PKS Pfur9-3_011125 has this domain architecture and groups phylogenetically with the atranorin cluster of *Cladonia rangiferina*. This cluster is present in both chemotypes and has a gene composition similar to the atr1 cluster of *C. rangiferina*, i.e., it has a *CytP450*, an *OMT* and a transporter gene. We propose that this cluster is most likely the atranorin cluster of *P. furfuracea*.

### The two genes for a metabolite FAS are present in *P. furfuracea*

*Aspergillus nidulans* has a metabolite FAS with properties similar to those expected for the unexplored lichen metabolite FASs. The *A. nidulans* FAS comprises two subunits, HexA and HexB, which produce and deliver to the aflatoxin NR-PKS the hexanoyl starter for norsolorinic acid, the first metabolite in the pathway (Brown *et al*., 1996; Watanabe and Townsend, 2002). We used the *Cladonia grayi* homologs of *HexA* and *HexB* (DalGrande et al., in preparation) to search for the corresponding genes in *P. furfuracea*. We found one 5619-bp HexA homolog and one 6285-bp HexB homolog. Like in *A. nidulans* and *C. grayi*, in both chemotypes of *P. furfuracea* these genes are adjacent and divergently transcribed from the same control region (genes FUN_005930 and FUN_005931 in the olivetoric acid chemotype; genes FUN_004275 and FUN_004276 in the physodic acid chemotype). These FAS subunit genes are not linked to the olivetoric/physodic cluster.

### Transcription of the olivetoric/physodic BGC and of *HexA* and *HexB*

We checked the transcription of the genes of interest (genes of cluster 4 and *HexA* and *HexB*) in lichen thalli of both chemotypes. In general, average transcription across the genome was lower in the olivetoric than in the physodic chemotype. This was reflected in the clusters as well. Transcriptome data suggest that in cluster 4, three genes out of 10, namely the *Pfur33-1_006185*, the *CytP450* and *gene6* (coding for an unknown protein) were transcriptionally active. We inferred the relative transcription activity by comparing the number of transcriptome raw reads (normalized by counts per million) that aligned to the respective gene (Table 4). Relative to the physodic acid (depsidone) chemotype, in the olivetoric acid (depside) chemotype all genes of cluster4 showed low transcription activity, although as compared to the other genes in the cluster the same three genes, *NR-PKS, CytP450* and *gene6* showed higher transcription activity. *HexA* and *HexB* were transcribed in both chemotypes. The number of read counts however, was higher in the physodic acid chemotype than in the olivetoric acid chemotype. This parallels the behavior of the cluster 4 genes (Table 4).

**Table 4.** Read count of the genes of cluster 4 and *P. furfuracea* homologs of *HexA* and *HexB* of the olivetoric- and physodic acid chemotype. Transcriptome raw reads and normalized read counts (by number of reads and gene length (RPKM approach)) aligned to the ten genes of the candidate physodic and olivetoric acid cluster. Genes in bold (gene 4, 5, and 6) are the ones with highest number of read counts.

| Gene | gene identity | gene length | physodic aid chemotype |  | olivetoric acid chemotype |  |
| --- | --- | --- | --- | --- | --- | --- |
|  |  |  | # raw reads aligned | reads (normalized by CPM) | # raw reads aligned | reads (normalized by CPM) |
| cluster 4 gene1 | unidentified | 3822 | 38 | 0 | no hits | - |
| cluster 4 gene2 | unidentified | 306 | no hits | - | 1 | 0 |
| cluster 4 gene3 | unidentified | 2685 | no hits | - | 19 | 0 |
| **cluster 4 gene4** | **cyt p450** | **2139** | **8677** | **117.29** | **1222** | **17.4** |
| **cluster 4 gene5** | **NR-PKS** | **6294** | **111226** | **511** | **1265** | **6.33** |
| **cluster 4 gene6** | **unidentified** | **2550** | **13690** | **155.25** | **6304** | **75.38** |
| cluster 4 gene7 | unidentified | 3284 | 1035 | 9.11 | 118 | 1.125 |
| cluster 4 gene8 | unidentified | 2191 | 936 | 12.35 | 205 | 1.98 |
| cluster 4 gene9 | unidentified | 492 | no hits | - | 12 | 0.73 |
| cluster 4 gene10 | monooxygenase | 1851 | 46 | 0.71 | 19 | 0.307 |
| FAS | FAS-A | 5619 | 3143 | **16.17** | 944 | **5.12** |
| FAS | FAS-B | 6285 | 1971 | **9.06** | 911 | **4.42** |

## Discussion

In this study we describe the putative BGCs for olivetoric- and physodic acid synthesis in the lichen-forming fungus *P. furfuracea* from high quality long-read genome assemblies of the two chemotypes. Furthermore, we identify the *HexA* and *HexB* homologs in *P. furfuracea*, likely to deliver the starters to the orcinol compound PKSs. Combining our findings with those of Kealey et al. (2021) and of other literature data, we propose an outline for the origin of the starter unit, chemotype variation, and synthesis of orcinol depsides and depsidones in lichens.

### True intraspecific variation underlies differences in BGCs between chemotypes

While most (51) BCGs were present in both chemotypes, six BGCs were present only in one chemotype (Supplementary info S2). In principle, the differences might be attributed to i) random variation in sequencing depth, ii) contamination by another fungus, and iii) true intraspecific variation. Random sequencing depth variation can be excluded, because we detected no reads of the missing BGC in the raw data. It is very unlikely that large genomic regions, such as entire BGCs, would be missed due to uneven coverage. Contamination from reads of minority fungal genomes (e.g. from lichenicolous fungi) can also be excluded, as we did not detect any coverage variation with underrepresented sequences compared to those from the main mycobiont. Furthermore, the clusters detected only in the physodic acid chemotype were also absent in the previously sequenced genome of *Pseudevernia furfuracea*, which was from the olivetoric acid chemotype (Meiser *et al*., 2017). True intraspecific variation is therefore the most likely cause of the observed differences in BGC content between chemotypes.

Intraspecies variations in BGCs have been reported for plants, bacteria, and fungi, and have been linked to ecological adaptation (Moore *et al*., 2014; Zhu *et al*., 2016; Thynne *et al*., 2019; Drott *et al*., 2020). For instance, the number of BGCs may vary between populations inhabiting different climatic conditions (Drott *et al*., 2020; Singh *et al*., 2021). In fact, fungal BGCs are suggested to be hotspots of gene gain/loss and duplication (Wisecaver *et al*., 2014; Lind *et al*., 2017; Rokas *et al*., 2018). Different strains of a single species can contain up to 15 strain-specific clusters (Vicente *et al*., 2018). The presence of unique BGCs suggests that each chemotype has a specialized metabolite potential based on genetic differences. Genome sequences of only two individuals are not likely to capture the pangenomic variation of BGCs within *P. furfuracea*. BGC variation among individuals of a species appears to be a common phenomenon and therefore a single individual may not represent the entire biosynthetic potential of a species (Susca *et al*., 2016; Villani *et al*., 2019; Singh *et al*., 2021). However, intraspecific biosynthetic variation can also arise when the BGCs involved are present in all individuals, as exemplified by the BGC likely responsible for the synthesis of olivetoric acid and physodic acid (see below).

Our results suggest that differences in presence/absence of BGCs are not linked to differences between chemotypes. Although here are many cluster differences between chemotypes, these differences do not affect the chemistry of the two chemotypes. Instead, the chemotypic differences appear to be because of the divergent regulation of the same cluster present in both chemotypes.

### The same candidate BGC is linked to depside and depsidone biosynthesis

The cluster we identified as the likely BGC linked to olivetoric/physodic biosynthesis has identical gene content in both chemotypes (Fig. 3), prompting the question of how one chemotype produces largely the depside olivetoric acid and the other largely the depsidone physodic acid. The likely BGC includes a *NR-PKS* and a *CytP450*, the two essential requirements for depside and depsidone synthesis (Armaleo et al. 2011; Kealey et al. 2021). These are also two of the three most highly transcribed genes in the cluster (Table 4). The function of the other genes, which are unidentified and mostly transcriptionally silent, with regard to olivetoric/physodic acid synthesis is unknown (Table 4). Theoretically, differential transcription of *CytP450* could explain the difference between chemotypes: while the *PKS* should be expressed in both chemotypes, repression of the *CytP450* gene in the olivetoric chemotype would prevent the depside to depsidone transition, whereas expression of the *CytP450* in the physodic chemotype would allow depsidone synthesis. However, the transcriptome data (Table 4) shows that *CytP450* is transcribed in both chemotypes, probably because averaging reads from thalli comprising different developmental and physiological stages cannot reflect subtle developmental transitions occurring at different times and locations. In fact each chemotype may occasionally produce both, the depside and the depsidone, depending upon the regulatory and other factors, but one of the two compounds remains below the level of detection. There are reports of occasional thalli of *P. furfuracea* containing both, physodic acid and olivetoric acid (Culberson *et al*., 1977).

While at a cellular scale differential transcription is decisive in determining phenotypes in fungi, secondary metabolite synthesis is a complex, multi-step process involving various genetic, epigenetic and environmental factors that together determine the spatio-temporal secondary metabolite profile of an organism (Fox and Howlett, 2008; Macheleidt *et al*., 2016; Keller, 2019). Often, the same BGC can be differentially regulated at the intraspecies level epigentically, posttranscriptionally or posttranslationally, to produce different compounds (Yin and Keller, 2011; Patra *et al*., 2013; Collemare and Seidl, 2019; Drott *et al*., 2020). For instance, the aspyridone cluster in *Aspergillus nidulans* can produce up to eight different compounds depending on the combination of genes activated (Wasil *et al*., 2013). Although our findings cannot explain which aspects of this complexity differentiate the chemotypes of *P. furfuracea*, they clearly indicate that differential regulation of the same BGC is involved. Our study shows that biosynthetic capabilities of organisms may vary within a species and highlights the importance of exploring the biosynthetic potential of organisms at the intraspecies level.

### A metabolite fatty acyl synthase is the likely provider of the hexanoyl starter for olivetoric acid synthesis

We found the homologs of *HexA* and *HexB* in *P. furfuracea*. As in *Aspergillus nidulans* and *Cladonia grayi* these genes are located next to each other and in divergent orientaion, suggesting that they are co-regulated by the same promoter. HexA and HexB refer respectively to the α and β subunits of the hexanoate synthase in *A. nidulans*. The FAS domains ACP, KR and KS are present in the α-chain and AT, ER, DH and malonyl-ACP transferase (MPT) in the β-chain (Jenni *et al*., 2006). HexA/B provides the hexanoyl starter to the PKS synthesizing the norsolorinic acid precursor of aflatoxin in *Aspergillus*. We propose that the HexA/B homolog in *P. furfuracea* delivers hexanoyl starters to initiate both rings of olivetoric acid, although the A-ring side chain ends up being two-carbons longer than the B-ring chain (Fig. 4), as described in the next section.

**Figure 4.**
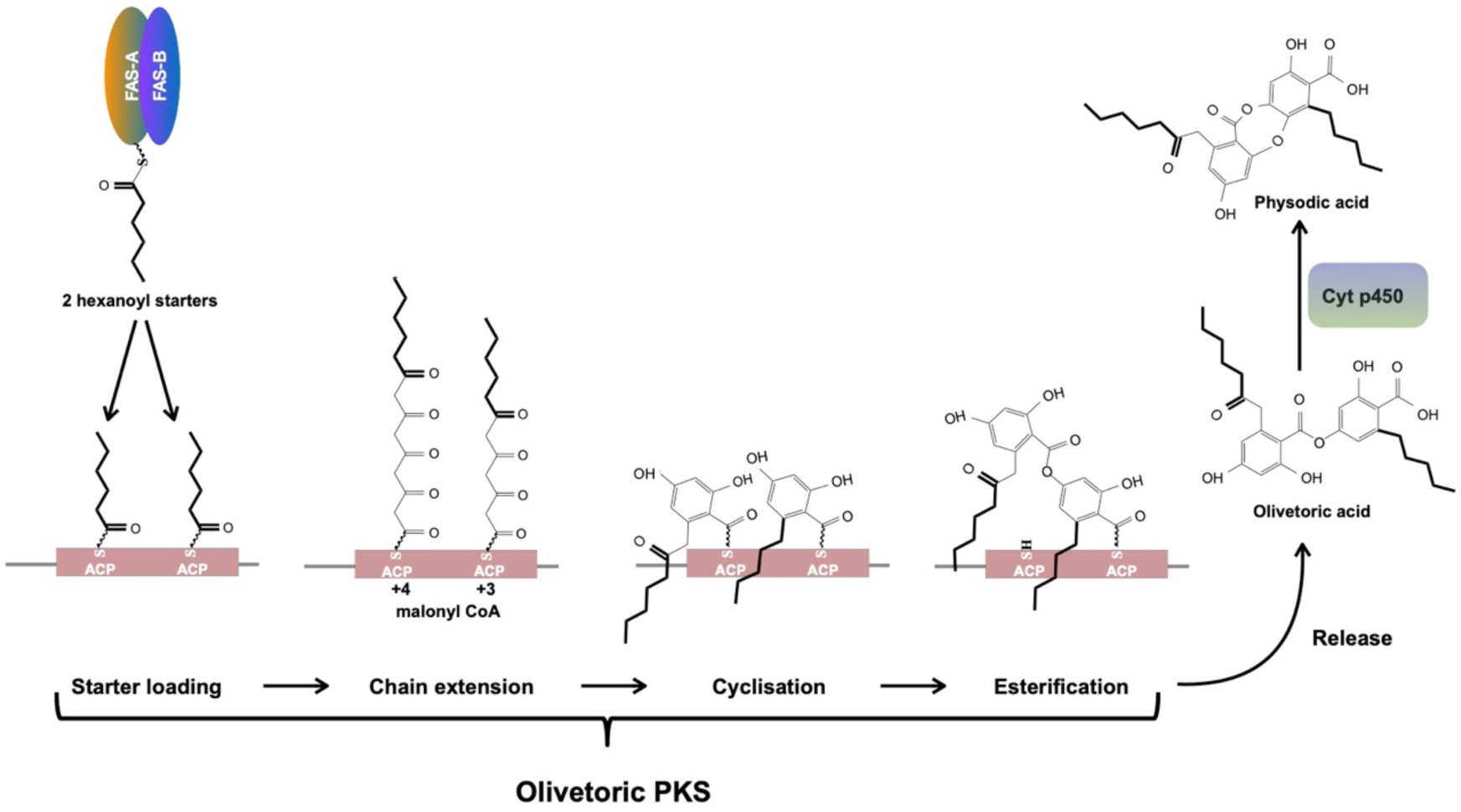
Putative scheme for depside/depsidone synthesis and function of FASs.

The *PKS* of cluster4 we identified in this study as most likely associated with olivetoric/physodic acid biosynthesis, is the same as *Pfur33-1_006185* that was heterologously expressed in yeast by Kealey et al. (2021). Interestingly, expression of this PKS in the heterologous host yielded lecanoric acid (Kealey et al. 2021), a compound never reported from thalli of *P. furfuracea*. Lecanoric acid and olivetoric acid differ in their starter side chains: both rings of olivetoric acid are started by an hexanoyl chain (Fig. 4) whereas both rings in lecanoric acid are started by the two-carbon chain from Acetyl CoA. This indicates that a PKS specific for olivetoric acid in *P. furfuracea* accepts Acetyl CoA as starter in yeast while it never does so in the lichen where it only accepts hexanoyl chains. Moreover, the heterologously expressed PKS continued to prefer acetyl CoA in yeast and produce lecanoric acid even when hexanoyl CoA was provided (Kealey et al. 2021). A likely solution to these apparent contradictions is that the “default” setting for the olivetoric PKS and perhaps for other orcinol depside PKSs is to accept free acetyl CoA as starter, but not free acyl CoAs with longer chains. This default setting is revealed only in the absence of a dedicated metabolite FAS like the one we identified in the lichen. Yeast has no metabolite FAS genes. The task of the metabolite FAS is to transfer directly to the PKS, through specific binding of the two proteins, the hexanoyl chain from the FAS ACP to the PKS ACP, with no free acyl CoA intermediate. That would explain why in yeast the olivetoric PKS would not use free hexanoyl CoA. Such an acyl-transfer mechanism is identical to what has been proposed for hexanoyl transfer between the HexA-HexB FAS in *A. nidulans* and the norsolorinic acid PKS (Watanabe *et al*., 1996; Watanabe and Townsend, 2002). Our identification in *P. furfuracea* of an expressed (Table 4) close homolog of the *A. nidulans* HexA-HeB FAS provides strong support for our proposed scenario. An important corollary of this scenario is that the side chain specificity in lichen orcinol compounds is not controlled exclusively by PKSs but likely results from specific protein-protein interactions between each PKS and a dedicated metabolite FAS which synthesizes and delivers the appropriate acyl-ACP starter directly to the PKS.

### An updated scheme of orcinol depside and depsidone synthesis

We combine here our results with those of Watanabe and Townsend (2002) Armaleo et al (2011), Feng et al. (2019), Lünne *et al*., (2020), and Kealey et al., (2021), to provide an updated scheme of orcinol depside and depsidone synthesis, using as example the synthesis of olivetoric and physodic acid. We limit our description to orcinol lichen compounds as we do not yet know how many of the same rules apply to ß-orcinol depsides and depsidones. Orcinol and ß-orcinol PKSs are separated by a deep evolutionary gulf (Fig. 2) and the biological differences between these two groups of lichen compounds are not well understood.

The scheme is depicted in Figure 4. Unless acetyl CoA provides the starter, as is the case for lecanoric acid and other orcinol compounds with methyl groups as side chains, a dedicated HexA/B FAS is needed to provide an acyl-ACP as starter to the *PKS*. In the case of olivetoric acid, hexanoyl-ACP is the starter for both rings and is transferred within the two proteins bound to each other from the FAS-ACP to the PKS ACPs (Fig. 4). The symmetric addition of starters is not the rule, as many orcinol compounds use different acyl chain starters for the two rings. Polyketide extension involves a minimum of three malonyl CoA additions, but can involve four. The PKS then cyclizes both polyketide chains to orcinol rings, esterifies the carboxyl of the A ring with the 4’ OH of the B ring and finally releases the depside by hydrolysis of the B ring thioester. The rings produced after four malonyl additions commonly have side chains with a ß-keto group derived from the carbonyl oxygen of the hexanoyl starter (Fig. 4), as seen on the A ring side chain of olivetoric and physodic acid. If the released depside is to be turned into a depsidone, a dedicated cytochrome P450 adds an ether bond, oxidatively coupling the C2 OH of the A ring to the 5’ C of the B ring.

## Conclusions

Our study contributes to the understanding of natural product synthesis in lichenized fungi in several ways. We identified the BGCs of the two *P. furfuracea* chemotypes and highlighted the putative cluster linked to physodic- and olivetoric acid biosynthesis. Additionally, we characterized the *P. furfuracea* homologs of *HexA*/*HexB*, the first FASs from lichen-forming fungi putatively involved in metabolite synthesis. Taken together, our results show that the same BGC has the potential to produce different compounds and suggests that intraspecific variation in the regulation of metabolite synthesis adds to the biosynthetic diversity and potential of organisms despite similar BGC content. Our study helped clarify some of the components determining chemotype variability in lichens and, in combination with other data, has allowed us to devise the most detailed scheme to date for the synthesis of orcinol depsides and depsidones. However, although the scheme combines the available evidence in a way consistent with the known molecular biology and biochemistry of these compounds, a number of details remain hypothetical and need experimental confirmation.

## Acknowledgements

We thank the LOEWE-Centre TBG funded by the Hessen State Ministry of Higher Education, Research and the Arts (HMWK). We thank Jürgen Otte (Frankfurt) for help in the lab, and Anjuli Calchera (Frankfurt) for HPLC analysis and technical support. The authors have no conflict of interest.

## Supporting information

**Table S1**. Voucher information of the samples used for the study

**Table S2**. Details of the estimated biosynthetic gene clusters in the two chemotypes of *P. furfuracea*. PKS number (column 2) refers to the most closely related PKS in the maximum likelihood tree presented in Figure 2. Region (column 3) refers to the cluster number and regions in the antiSMASH output.

